# Seed biopriming with P- and K-solubilizing *Enterobacter hormaechei* sp. improves the early vegetative growth and the P and K uptake of okra (*Abelmoschus esculentus*) seedling

**DOI:** 10.1101/2020.04.24.059303

**Authors:** Muhamad Aidilfitri Mohamad Roslan, Nurzulaikha Nadiah Zulkifli, Zulfazli M. Sobri, Ali Tan Kee Zuan, Sim Choon Cheak, Nor Aini Abdul Rahman

## Abstract

Limited information is available that seed biopriming by plant growth-promoting bacteria such as those among *Enterobacter* spp. play a prominent role to enhance vegetative growth of plants. Contrary to *Enterobacter cloacae*, *Enterobacter hormaechei* is a less-studied counterpart despite its vast potential in plant growth-promotion mainly through the inorganic phosphorus (P) and potassium (K) solubilization abilities. To this end, 18 locally isolated bacterial pure cultures screened and three strains showed high P- and K-solubilizing capabilities. Light microscopy, biochemical tests and 16S rRNA gene sequencing revealed that strains 15a1 and 40a were closely related to *Enterobacter hormaechei* while strain 38 was closely related to *Enterobacter cloacae* (Accession number: MN294583; MN294585; MN294584). All *Enterobacter* spp. shared common plant growth-promoting traits, namely N_2_ fixators, indole-3-acetic acid producers and siderophore producers. Gibberellic acid was only produced by strain 38 and 40a, while exopolysaccharide formation was solely detected on agar containing colonies of strain 38. Under *in vitro* germination assay of okra (*Abelmoschus esculentus*) seeds, *Enterobacter* spp. significantly improved overall germination parameters and vigor index (19.6%) of seedlings. The efficacy of root colonization of *Enterobacter* spp. on the pre-treated seedling root tips was confirmed using Scanning Electron Microscopy (SEM). The pot experiment of bioprimed seeds of okra seedling showed significant improvement of the plant growth (> 28%) which corresponded to the increase of P and K uptakes (> 89%) as compared to the uninoculated control plants. The leaf surface area and the SPAD chlorophyll index of bioprimed plants were increased up to 29% and 9% respectively. This report revealed that the under-explored species of P- and K-solubilizing *Enterobacter hormaechei* sp. with multiple plant beneficial traits hold as a good potential sustainable approach for enhancement of soil fertility and P and K uptakes of plants.

## Introduction

The application of sizable inputs of synthetic chemical fertilizer such as those containing inorganic phosphorus (P) and potassium (K), have long been employed in the conventional agriculture practices. While the outcome of this approach has been an inextricable success, the excessive application of chemical fertilizer particularly by large scale farming has posed detrimental impacts on environmental quality and economic growth. The plausible long-term risks and threats inflicted by the profuse usage of chemical fertilizer are still limited [1]. However, there have been several reported cases of groundwater and waterways contamination due to nutrient leaching of P, K and Nitrogen (N_2_), especially in the areas which received high rainfalls and near streams or drains [2–5]. Such environmental issues have been lengthily discussed as to mitigate the loss of nutrients added to the soil and the risk of eutrophication occurring in the adjacent water bodies. Therefore, the exploration of alternative strategies that can promote competitive crop growth while maintaining the environmental safety is definitely a timely measure.

Recently, biofertilizer application has shown a promising practice for sustainable production of agricultural yields while imposing cutbacks up to 50% on the required chemical fertilizer dose into the soils [1,6,7]. The term biofertilizer or interchangeably called as bioinoculant or plant growth-promoting (PGP) microorganisms is described as a substance which comprises beneficial living microorganisms as seed, plant or soil application to promote plant growth by elevating the supply or availability of host-specific essential nutrients [8]. Plant growth-promoting bacteria (PGPB) for instance, may benefit plant growth typically by either: facilitating the acquisition of soil nutrient by asymbiotic N_2_-fixation and inorganic phosphorus (P) and potassium (K) solubilization; modulating the level of phytohormone such as indole-acetic acid (IAA), cytokinins, and gibberellins; resisting against plant-pathogenic microorganisms [1].

At present, majority of field-tested bacterial taxonomic groups affiliated to PGPB belonged to ranges of genera: *Azospirillum, Rhizobium, Agrobacterium, Rhizobium* (Alphaproteobacteria); *Azoarcus, Burkholderia*, (Betaproteobacteria); *Acinetobacter, Enterobacter, Klebsiella, Pantoea, Pseudomonas, Serratia*, (Gammaproteobacteria); *Bacillus, Frankia* (gram-positive bacteria) [7,9–11]. *Enterobacter cloacae* for instance, is a well-studied species concerning its immense plant beneficial effects on a wide array of crop varieties. The efficiency of endophytic *E. cloacae’s* application on crop yield productivity have been discussed in major details along with its functional phenotypic, genotypic and metabolomic roles behind it [6,12–14]. Researchers, lately, have started to explore the PGP potential of its counterpart, *Enterobacter hormaechei*, which have been reported to serve as biological fungicide against *Fusarium* spp. [15] and improved the plant biomass and biosynthesis of aloin-A in aloe vera [16]. However, information that is available concerning the PGP potential of *E. hormaechei* is scarce and further investigation is required to understand the biological activities and soil-plant-interactions involved.

While there are myriads of techniques to introduce PGPB to crops, seed biopriming is one of the preferable methods to perform effective seed surface bacterial inoculation. Biopriming is conceptualized as a technique of seed priming using living bacterial inoculum which allows the bacterial adherence and acclimatization to the seeds in the prevalent conditions [17]. Seed biopriming provides a lot of benefits to plants especially in enhancing seed viability, germination, seed vigor, growth, and yield [18]. In fact, bioprimed seeds have been proven to significantly enhance the germination percentage and the germination rate of seeds even under an induced environmental stress circumstance such as osmotic stress condition [19].

Keeping in view the importance of growth-limiting nutrient for plants such as P and K, the present study was designed to screen and characterize P- and K-solubilizing bacteria from our bacterial culture collection, retrieved from our previous bacterial isolation work on oil palm empty fruit bunch (OPEFB) and chicken manure co-composting process [20]. Additionally, the selected strains were screened for various PGP properties such as N_2_ fixation, indole-3-acetic acid production (IAA), gibberellic acid (GA3) production, zinc solubilization, siderophore and exopolysaccharide (EPS) production. To assess the effects of seed biopriming with the selected strains on okra seedling growth performance, seed germination bioassay and pot experiment under net house condition were performed thereafter.

## Materials and methods

### Preliminary assay of qualitative P and K solubilization

Eighteen bacterial isolates were revived from glycerol stock stored in –80°C freezer in Research Laboratory 1.6, Bioprocessing and Biomanufacturing Research Center, Faculty of Biotechnology and Biomolecular Sciences, Universiti Putra Malaysia. The isolates were previously isolated from several sources, such as from agricultural soil, oil palm empty fruit bunch (OPEFB) and chicken manure co-composting process [20]. Isolates were grown on nutrient agar at 30°C for 24 h and routinely subcultured prior to every assay.

Phosphate solubilization activity of isolates was evaluated using National Botanical Research Institute’s Phosphate growth medium (NBRIP) which contains (per Liter): 10 g glucose; 5 g Ca_3_(PO_4_)_2_; 5 g MgCl_2_·6H_2_O; 0.25 g MgSO_4_·7H_2_O; 0.2 g KCl, 0.1 g (NH_4_)_2_SO_4_ and 15 g agar [21]. A modified NBRIP medium was prepared separately by addition of pH indicator of 10 mL bromothymol blue which contained 0.5% aqueous solution dissolved in 0.2 N KOH [22]. Plate assay for potassium solubilization was performed using Aleksandrov agar (Himedia) containing insoluble potassium aluminosilicate. A modified Aleksandrov agar was prepared separately with the addition of 0.018 g/L phenol red dye as a pH change indicator [23]. A fresh 24 h colony culture was spot on the solidified agar using a sterile inoculating needle and the plate was incubated at 30°C for 5 days in triplicate. The halo zone formation was interpreted as solubility index (SI) which from now onwards was termed as P solubility index (PSI) and K solubility index (KSI) respectively and determined using the the ratio of the total halo diameter to the colony diameter [6].

### Antagonistic interaction assay

Antagonistic interaction assay between selected isolates was characterized by pour plate method as described by Grossart et al., (2004). A molten nutrient 1% agar (2.5 mL) was mixed with 50 μL target isolate suspension (1×10^8^ CFU/mL). The cell suspension agar was poured onto a nutrient agar plate (Sigma) and left for 10 min until it was completely solidified. An aliquot (10 μL) of test strain (1×10^8^ CFU/mL) was applied onto the lawn and thereafter incubated at 30°C for 3 days. The strains were tested in triplicate against each other and observed daily for inhibition zones. Inhibitory activity was recorded when the inhibition zone was equal to or more than 4 mm of the diameter of the applied colony in both parallels.

### Determination of quantitative P and K solubilization in liquid culture

Another experiment was set up to estimate the quantitative P and K solubilization of selected isolates in liquid media. Bacterial cultures (inoculum adjusted to 1×10^8^ CFU/mL) were transferred to NBRIP and Aleksandrov liquid media respectively. For a synergistic interaction test, a mixed bacterial culture was prepared by mixing inoculum culture at equal ratio 1:1:1 v/v of every isolate prior to inoculation into test media. Cultures were grown at 30°C for 5 days on a rotary shaker at 150 rpm. The pH change of the liquid media was measured at the end of incubation time and cultures were centrifuged at 10,000 rpm for 10 min to obtain cell-free supernatant. The P solubilization in liquid media was determined by yellow phospho-molybdo-vanadate colorimetric method [25] using UV-VIS spectrophotometer Secomam Uviline 9400 (France). The concentration of soluble P was determined according to the dilution series of standard P graph plot of KH_2_PO_4_ stock solution (50 μg/mL P). The K solubilization was determined by atomic absorption spectrometer (AAS) using flame air-C_2_H_2_ at the wavelength of 766.5 nm [1,26].

### Organic acid determination in post-incubation liquid culture

The cell-free supernatant of NBRIP and Aleksandrov liquid culture was filtered using 0.22 μm nylon filter (Millipore, USA) respectively. About 20 μL of cell-free supernatant was passed through Rezex (Phenomenex) organic acid (ROA) column in Agilent high performance liquid chromatography (HPLC) system at 60°C with flow rate of 0.6 mL/min. Organic acid standard solutions were prepared beforehand which were citric, gluconic, acetic, GA3, malic, succinic and lactic acid at 10 mg/mL concentration. Quantitative estimation of organic acids was performed based on the comparison of peak area and retention time of sample with those of standards.

### Bacterial growth curve, biochemical tests and molecular identification

The selected bacteria were identified by phenotypic and genotypic methods. The growth curve pattern of bacteria was determined using nutrient broth (Sigma) which received standardized amount of 1 mL starting inoculum (1 × 10^8^ CFU/mL) in 250 mL Erlenmeyer flask. The culture was incubated at 37°C on a rotary shaker at 150 rpm and thereafter sampled (2 mL) every 2 h. At the time of sampling, absorbance reading at 600 nm was recorded using UV/VIS spectrophotometer. In another experiment, bacteria were screened for a series of biochemical reaction tests: amylase [27]; [28]; protease [29]; lipase [30]; catalase; urease; acetate utilization. Olympus CH light microscope (Tokyo, Japan) was used to observe morphological characteristics and Gram’s reaction of the bacterial isolates. Bacterial 16S rRNA gene was analyzed by employing DNA barcoding service at Apical Scientific Laboratory, Selangor, Malaysia, using BigDye^®^ Terminator v3.1 cycle sequencing kit and sequenced by Applied Biosystems genetic analyzer platform.

The generated sequences were viewed and trimmed using BioEdit version 7 [31] and compared with the closest strains in the GenBank database using Basic Local Alignment Search Tool (BLAST) in terms of percent identity, query coverage and E-value. The phylogenetic tree was reconstructed using MEGA X Alignment Explorer [32] as sequences were aligned by ClustalW and inferred by using Neighbour-Joining method with 1000 replicates of bootstrap values (Saitou and Nei, 1987). Sequences were deposited into National Center for Biotechnology Information (NCBI) GenBank database via online sequence submission tool BankIt and the sequence accession numbers were retrieved.

### Plant growth-promoting activity assays

The *Enterobacter* spp. were subjected to further screening for PGP activities e.g., N_2_ fixation, IAA production, GA3 production, zinc solubilization, siderophore and EPS production. The quantitative estimation of nitrogen fixation activity was performed using a modified micro Kjedahl method [34]. Cultures were grown in N-free malate semisolid medium (NFM) [35] at 30°C for ten days on a rotary shaker at 150 rpm. The change of colour of the light green NFM to blue indicates the ability of isolates to convert atmospheric N_2_ into ammonia. The percentage of fixed nitrogen was measured using sulfur digestion and distillation with 10 mol/L NaOH [36].

Tryptic soy broth was employed to evaluate the IAA production of *Enterobacter* spp. with the addition of 0.1 g/L L-Tryptophan as the precursor of IAA in the culture composition [37]. Cell-free supernatant was mixed with 2 mL Salkowsky reagent containing 2% of 0.5 M FeCl_3_ in 35% perchloric acid [38] and was allowed to settle for 25 min. The intensity of pink color was determined using UV spectrophotometer at 535 nm and compared against the standard curve of pure IAA of known concentration.

The *Enterobacter* spp. were screened for zinc solubilization by using tris-minimal salt media which contains (per Liter): 10 g D-glucose, 6.06 g Tris-HCl, 4.68 g NaCl, 1.49 g KCl, 1.07 g NH_4_Cl, 0.43 g Na_2_SO_4_, 0.2 g MgCl_2_.2H_2_O, 0.03 g CaCl_2_.2H_2_O, and 15 g agar. The ZnO (1 g/L) was added in the media as the sole source of zinc to check the ability of isolates to solubilize zinc oxide [39]. The plate was incubated at 30°C for 5 days in triplicate and formation of halo zone around colonies indicated zinc solubilization activity. Zinc solubilization index (ZSI) was calculated as the ratio of the total halo diameter to the colony diameter [6].

The siderophore production of *Enterobacter* spp. was assessed qualitatively as described by Lakshmanan et al., (2015) according to Schwyn and Neilands (1987) protocol using chrome azurol S (CAS). The reagent was prepared by dissolving 0.0605 g CAS in 50 mL water and mixed with 10 mL FeCl_3_ solution (1 mM FeCl_3_.6H_2_0, 10 mM HCl). With constant stirring, the CAS-FeCl_3_ solution was slowly added into hexadecyltrimethylammonium solution (0.0729 g in 40 mL distilled water). The final mixture of 100 mL was added into 900 mL of autoclaved Luria broth (LB) agar, pH 6.8. Bacterial isolates exhibiting a yellowish-orange halo zone after 10 days of incubation were considered siderophore-producing strains.

EPS production of *Enterobacter* spp. was evaluated based on the formation of a mucoid colony on the agar plate-containing glucose such as the NBRIP and Aleksandrov agar after ten days of incubation. This biopolymer formation was verified by mixing a portion of the mucoid substance in 2 mL of absolute alcohol (99%). Formation of precipitate at the bottom of solution indicated EPS production [42].

### Biosafety inspection of *Enterobacter* spp

Biosafety inspection of *Enterobacter* spp. was carried out using blood agar medium to evaluate the haemolytic potential [43]. An overnight culture of *Enterobacter* spp. was streaked onto a blood agar plate comprising 5% (v/v) sheep blood and incubated at 30°C for 48 h. Haemolysis of red blood cells was observed by the formation of a clear zone or greenish colouration around colonies indicating *β*- or *α*-haemolysis respectively, while no clear zone indicates **γ**-haemolysis.

### Okra seed biopriming and germination bioassay

Okra variety seeds of F_1_ Hybrid Okra-Best Five 304 (Green World) were surface sterilized with 70% ethanol and diluted Clorox^®^ solution (3%) for 5 min and 5 sec respectively followed by triple washes with sterile distilled water. The sterilized seeds were dried in laminar flow with constant air flow at room temperature for 1 h and thereafter soaked in a freshly grown *Enterobacter* spp. culture (adjusted to 1×10^8^ CFU/mL) suspended in 0.85% phosphate buffered saline (PBS) separately for 1 h. Three seeds were placed into plant tissue culture tube containing 50 mL of 0.25% water agar [6] with the additional nutrient of Murashige and Skoog basal medium (4.4 g/L). Tubes were placed in sterile tissue culture room at 26±2°C for 3 days for water imbibition period in darkness, and subsequently continued for 4 days under controlled light condition (16 h light, 8 h darkness) for radicle protrusion period. Uninoculated seeds soaked in sterile PBS were used as control. The experiment was set up using completely randomized design (CRD) with six replicates for each treatment where each tube contained three seeds.

Several parameters were examined after 7 days post-inoculation (DPI) such as the length of hypocotyl and radicle, and the number of lateral roots formed. Measurement was taken from six seedlings sampled randomly from each treatment. Germination in seeds was achieved when radicals are half the size of the seed. Percentage (%) of germination and seedling vigor index were determined using the formula described by [44]. The experiment was repeated twice.

Root colonization potential of *Enterobacter* spp. on okra seedlings root tips was observed via electron microscopy. Root samples (1 cm) were prepared aseptically and placed into vials for each treatment. Glutaraldehyde (4%) was added into the vials to fix samples for 2 days at 4°C. Samples were washed three times with 0.1 M sodium cacodylate buffer of 30 min soaking each. Osmium tetroxide (1%) was used for post-fixation of roots for 2 h at 4°C prior to rewashing three times with 0.1 M sodium cacodylate buffer. Roots were dehydrated with a series of acetone washing of different concentration from 35% to 100% at almost 1 h incubation time each. Critical point dryer Autosamdri^®^-815 was used for final dehydration for about 1 h before mounting onto the stub. Dehydrated roots were coated with gold using sputter coater and transferred to slide for viewing using scanning electron microscopy (SEM) Jeol JSM-6400.

### *In planta* evaluation of seed biopriming with *Enterobacter* spp

P and K solubilizing *Enterobacter* spp. were evaluated for their plant growth promoting potential by pot experiment using Holland peat moss (pH 7.5, Available P 36.4 mg/Kg, Total P 648 mg/Kg, Available K 57.6 mg/Kg, Total K 1560 mg/Kg) under net house conditions. The experiment was performed at Research Laboratory 1.6, Bioprocessing and Biomanufacturing Research Center, Universiti Putra Malaysia (3°00’08.3”N 101°42’12.0”E) during the rainy season (November–December 2019). Okra variety seeds of F_1_ Hybrid Okra-Best Five 304 (Green World) were surface sterilized and bioprimed with *Enterobacter* spp. as described previously, except for uninoculated seeds where sterile PBS was used for priming. Carrier material of autoclaved peat moss (50 g per pot) was mixed with an overnight culture of *Enterobacter* spp. (1×10^8^ CFU/mL) and left for 1 h.

Inoculated seeds were sown in sowing pots with 8 cm diameter containing 0.2 Kg peat moss which was previously mixed with the prepared carrier. Pots were watered thoroughly after sowing to serve a good moisture condition for a favourable seed germination. There were three seeds per pot with six replicates under four different treatments and arranged in CRD. Seedlings were uprooted after 25 days of sowing and studied for plant growth parameters e.g., shoot length, root length, wet and dry biomass. Leaf surface area was measured using calculation described by Alami et al., (2018) while the chlorophyll index was determined using Chlorophyll Meter SPAD-502 Plus (Konica Minolta) according to the standard manual. The experiment was repeated twice successively. Soil viable cell was quantified by a standard serial dilution method using phosphate buffered saline (pH 6.8) and nutrient agar.

### P and K content determination of seedlings and soil

Peat soil as well as the leaf, stem and roots of the uprooted seedlings were dried at 70°C for 48 h until a constant weight was achieved. Samples were ground finely and weighed at 0.25 g separately. The dried ground samples were subjected to one-step concentrated nitric acid digestion method [46] for total P and K content estimation. Available P and K were determined by sodium carbonate extraction method [47]. The determination of P and K content was carried out by inductively coupled plasma optical emission spectrophotometer (ICP-OES) Optima 7300DV (Perkin Elmer) using P and K analytes of 213.617 nm and 766.49 nm respectively.

### Statistical analysis

Mean comparison of solubilization index, pH of culture media, soluble P and K, germination, seedling growth parameters and the P and K content were analyzed using one-way analysis of variance (ANOVA) by using SPSS software Package Version 25.0 (SPSS Inc., USA). Difference between treatments was compared by least significant difference (LSD) test at P<0.05.

## Results

### Preliminary screening of P and K solubilizing bacteria

All bacterial cultures were retrieved from glycerol stocks of our previous bacterial isolation works from several sources, such as from agricultural soil, oil palm empty fruit bunch (OPEFB) and chicken manure co-composting process. In total, 18 phenotypically different isolates were subcultured and strictly selected from those which were able to optimally solubilize inorganic P and K on solid NBRIP and aleksandrov agar respectively. Three out of eighteen bacterial isolates, labelled as 15a1, 38 and 40a (later were identified as *Enterobacter* spp.) exhibited the largest halo zone on agar media for both P and K solubilization activity respectively (**Table 1**). The PSI and KSI of the three isolates were determined in the next experiment.

**Table 1.** Qualitative P and K solubilization screening of *Enterobacter* spp.

| Isolate | P solubilization | K solubilization |
| --- | --- | --- |
| 15a1 | ++ | ++ |
| 38 | ++ | ++ |
| 40a | ++ | ++ |
| 1c | — | — |
| Sp4 | — | — |
| 2b | — | — |
| 2d55 | — | — |
| 10c | — | — |
| 12a | — | — |
| 14a | — | — |
| 15-1 | — | — |
| 15a2 | ++ | — |
| 16b35 | — | — |
| 20b | + | — |
| 22a | — | — |
| Sb16 | + | — |
| Psb16 | + | — |
| Bc | + | — |

Note: P and K solubilization were categorized as low activity when the halo zone is < 5 mm (denoted as +) or high activity when the halo zone > 5 mm (denoted as ++).

Based on *in vitro* plate screening (Fig 1), *E. hormaechei* 40a produced the highest PSI (2.5 on NBRIP, 2.8 on modified NBRIP) while *E. hormaechei* 15a1 surpassed *E. clocae* 38 and *E. hormaechei* 40a for KSI level (3.2 on Aleksandrov agar). However, *E. hormaechei* 40a showed the highest KSI (4.3) on modified Aleksandrov agar as compared to the other two strains. All of them showed a prominent size of the acidic zone on both modified NBRIP and Aleksandrov agar containing pH indicator dye with *E. hormaechei* 15a1 and 40a showing the largest zone (72.7 mm and 69.3 mm respectively on modified NBRIP).

**Fig 1.**
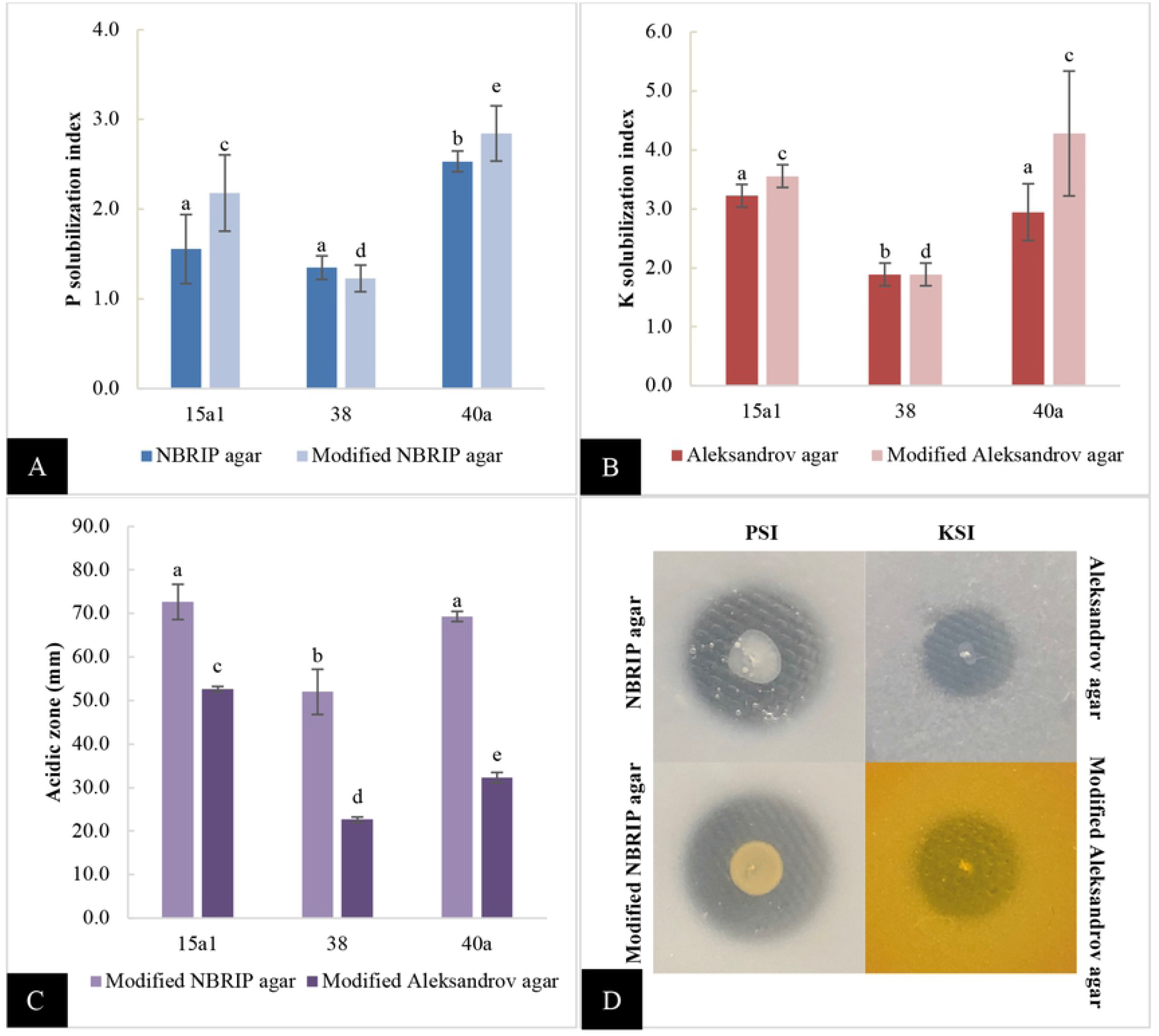
*In vitro* P and K solubilization index of *Enterobacter* spp. on different solid agars. A: PSI on NBRIP and modified NBRIP agar, B: KSI on Aleksandrov and modified Aleksandrov agar, C: acidic zone on modified NBRIP agar and modified Aleksandrov agar, D: halo formation due to solubilization of insoluble P and K on different agars. All values are the mean value of three biological replicates alongside the standard deviation indicated by the error bar. Means followed by different letters in the same category indicate significant differences at P<0.05.

A mixed culture of *Enterobacter* spp. was prepared to contain equal ratio of inoculum size as stated in the previous section. Based on plate assay, there were no observable inhibitory activities occurred during growth of bacteria against each other. It was presumed that the mixed culture could demonstrate synergistic activities when tested for the subsequent PGP trait assays. Thus, the following qualitative estimation of P and K solubilizing activities of *Enterobacter* spp. incorporated both individually and mixed culture experiments.

### Quantitative estimation of P and K solubilizing activity and organic acid production

In the quantitative assay of P solubilizing activity, *E. hormaechei* 40a consistently demonstrated the highest soluble P (508.25 μg/mL) followed by *E. clocae* 38 (471.58 μg/mL) and *E. hormaechei* 15a1 (450.75 μg/mL) in NBRIP media after 5-day incubation (Fig 2). Contrary to K solubilizing activity, *E. hormaechei* 15a1 released the highest soluble K (72.90 μg/mL) followed by *E. clocae* 38 (71.15 μg/mL) and *E. hormaechei* 40a (65.95 μg/mL) in Aleksandrov culture. The pH level of the inoculated cultures was significantly higher (P<0.05) than uninoculated control media (7.13 in NBRIP broth, 6.67 in Aleksandrov broth) and ranged between acidic pH of 3.78 (40a in Aleksandrov broth) to 4.85 (38 in Aleksandrov broth). However, the mixed culture did not show any synergistic effects on the solubilization of P and K as well as the pH level of the culture in comparison with those as single cultures (P>0.05).

**Fig 2.**
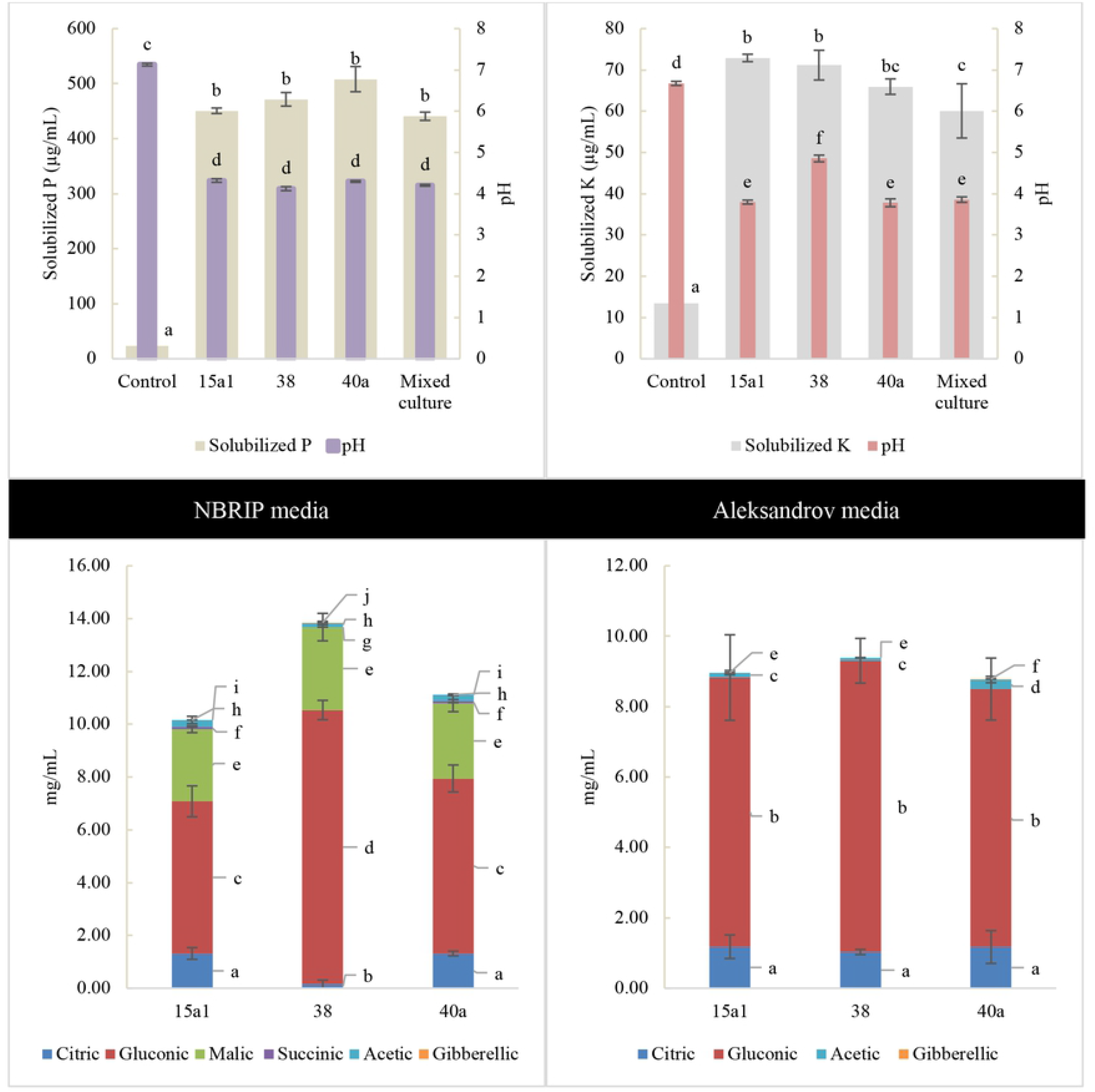
Interaction between pH level and solubilization of P and K in liquid NBRIP and Aleksandrov media respectively after 4^th^ of day of incubation. Organic acids produced by *Enterobacter* spp. were quantified using Agilent HPLC system against the known standard of citric, gluconic, malic, succinic, acetic, gibberellic and lactic acid respectively. All values are the mean value of three biological replicates alongside the standard deviation indicated by the error bar. Means followed by different letters in the same category indicate significant differences at P<0.05.

The production of organic acids by *Enterobacter* spp. were identified using HPLC and the resulting chromatogram profiles were analysed against the standards (S2 Fig). In NBRIP culture, all three bacteria produced a prominent amount of gluconic acid, malic acid and citric acid. *E. clocae* 38 produced the highest amount of gluconic acid (10.39 mg/mL) followed by malic acid (3.14 mg/mL) while both *E. hormaechei* 15a1 and 40a produced higher amount of citric acid (1.32 mg/mL) as compared to *E. clocae* 38. Succinic, acetic and GA3 acid were produced in a small amount (< 0.3 mg/mL) by *E. hormaechei* 15a1 and 40a. Likewise in Aleksandrov media, all *Enterobacter* spp. produced a prominent amount of gluconic and citric acid. Contrary to NBRIP media, a minimal amount (< 0.3 mg/mL) of acetic acid was detected in Aleksandrov media, while a consistent low amount of GA3 acid was detected in both media.

### Phenotypic and molecular identification of isolates

The growth curve of *Enterobacter* spp. was recorded by optical density (600 nm) using nutrient broth throughout 24-h incubation (Fig 3). The maximum cell density recorded for *E. hormaechei* 15a1 (2.46) was at 18^th^ h while the cell growth peak of *E. clocae* 38 and *E. hormaechei* 40a was recorded at 20^th^ h (2.23 and 2.53 respectively). Cell morphology observation under a light microscope revealed that cells were single coccobacillus and Gram negative. The bacterial colony was observed as round, white, entire, raised, butyrous and were around 0.8 to 1.5 mm diameter size on nutrient agar.

**Fig 3.**
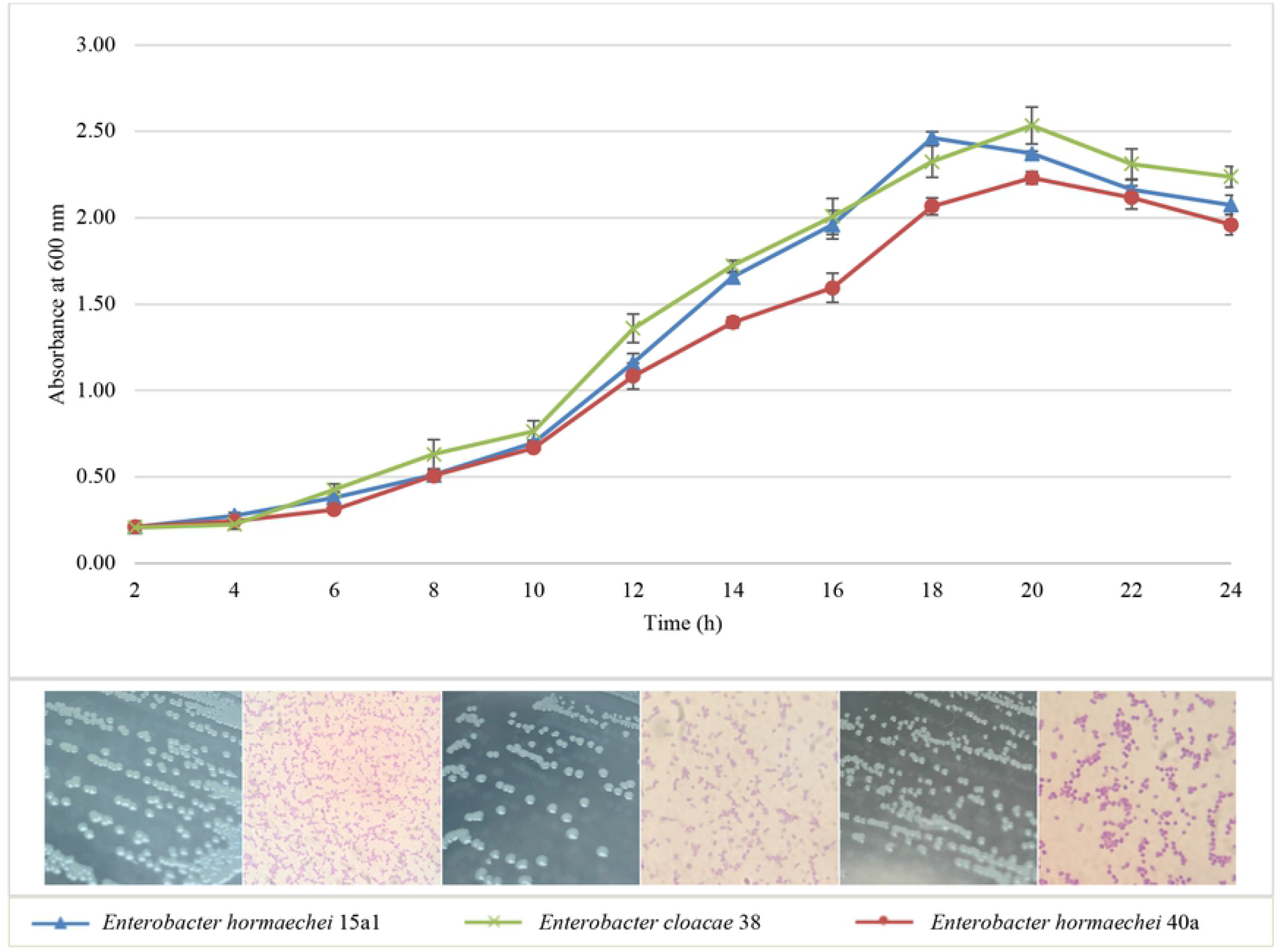
Growth of *Enterobacter* spp. in nutrient broth at 37°C shaking at 150 rpm in 24 h. The graph legend shows the colony morphology of *Enterobacter* spp. on nutrient agar and the cell morphology under light microscope at 1000× magnification.

Biochemical characterization assay of *Enterobacter* spp. showed that none of the strains exhibited hydrolytic activities such as amylase, cellulase, protease and lipase activities (Table 2). Catalase and urease assay were tested positive for all *Enterobacter* spp. while acetate utilization was solely demonstrated by *E. cloacae* 38. PCR amplification of 16S rRNA gene sequencing was employed to identify the P and K solubilizing strains. Sequences were compared on the NCBI BLASTN tool based on percentage similarity, E-value and query coverage (S1 Table). BLAST result of 15a1 and 40a presented 100% similarity with *Enterobacter hormaechei* whereas 38 showed 100% similarity with *Enterobacter cloacae*. Sequences were deposited to NCBI GenBank database and the unique accession numbers were retrieved (MN_2_94583.1 - MN_2_94585.1).

**Table 2.**
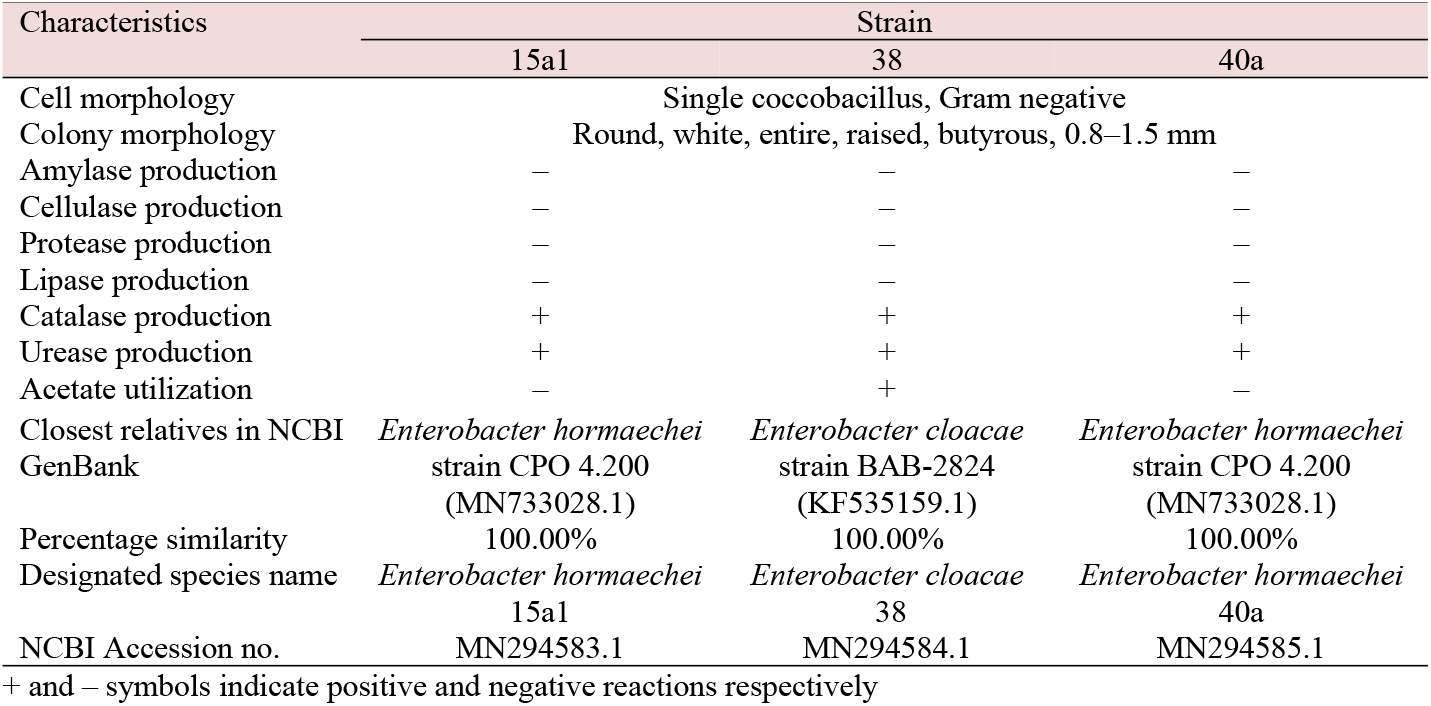
Cell and colony morphology, biochemical characteristics and molecular identification of *Enterobacter* spp.

The evolutionary relationship was reconstructed using the neighbour-joining method [33] to generate the optimal phylogenetic tree with the sum of branch length of 0.14724404 (Fig 4). The numbers shown next to the branches represented the replicate trees percentage in which the related taxa were grouped together in the bootstrap test of 1000 replicates [48]. p-distance was used to compute the evolutionary distances among strains [49] which in total involved 14 nucleotide sequences. Pairwise deletion was opted to eradicate ambiguous positions for each sequence pair. The final dataset consisted of 1506 positions and all evolutionary analyses were performed using MEGA X [32,50].

**Fig 4.**
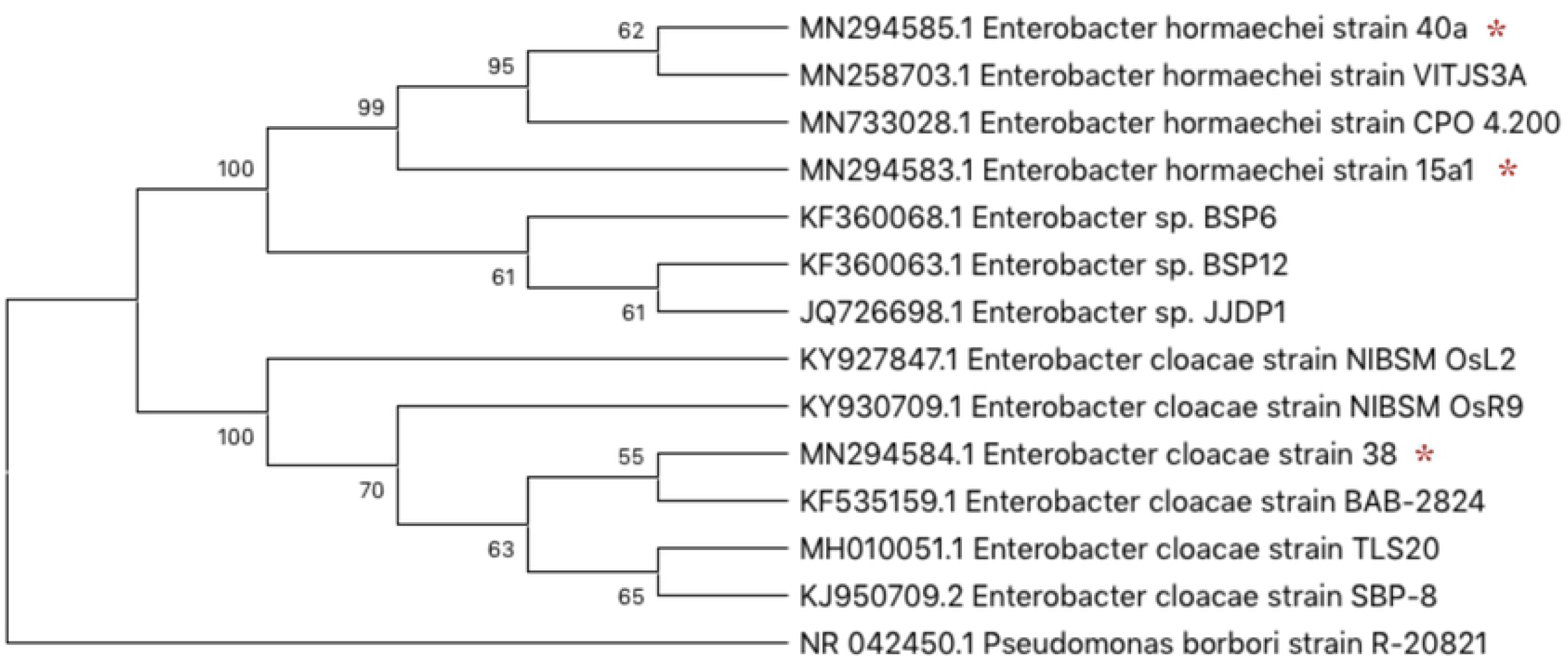
Molecular identification of *Enterobacter* spp. via 16S rRNA gene sequencing. Optimal phylogenetic tree of isolates (indicated with asterisk) and their closest strains was reconstructed by Neighbor-joining method with the sum of branch length of 0.14724404 using MEGA X Alignment Explorer.

### Multi-putative plant growth promoting activity of *Enterobacter* spp

*Enterobacter* spp. were also examined on the ability to demonstrate multi-putative plant growth promoting activities such as N_2_ fixation, IAA production, GA3 production, zinc solubilization, siderophore and exopolysaccharide (EPS) production (Table 3). N_2_ fixation of *Enterobacter* spp. was detected by the colour change of the light green NFM to blue indicating ammonia production from the conversion of atmospheric N_2_. It was later confirmed by the micro Kjedahl method that all strains were able to fix N_2_ whereby *E. cloacae* 38 fixed the highest N_2_ (2.10 μg N/mL) followed by *E. hormaechei* 15a1 (1.23 μg N/mL) and *E. hormaechei* 40a (0.36 μg N/mL).

**Table 3.** Multi-putative plant growth promoting characteristics of *Enterobacter* spp.

| Isolate | Fixed N<sub>2</sub> (µg N/mL) | IAA (µg/mL) | GA3 (µg/mL) | Zinc | Siderophore | EPS |
| --- | --- | --- | --- | --- | --- | --- |
| *E. hormaechei* 15a1 | 1.23±0.51 | 110.81±0.83 | NA | – | ++ | – |
| *E. cloacae* 38 | 2.10±0.82 | 184.03±0.93 | 21.05±0.51<sup>a</sup> | – | + | + |
| *E. hormaechei* 40a | 0.36±0.44 | 57.51±2.81 | 23.16±0.91<sup>b</sup> | – | ++ | – |

Gibberellic acid were detected using HPLC system in NBRIP <sup>a</sup> or/ and Aleksandrov <sup>b</sup> culture
Data are mean value of three biological replicates with ± denoting standard deviation. + and – indicates positive and negative traits respectively while ++ indicates high intensity qualitatively.

Additionally, *E. cloacae* 38 produced the highest IAA in LB-tryptophan media (184 μg/mL) compared to the other two strains. While in GA3 assay, both *E. cloacae* 38 and *E. hormaechei* 40a produced a detectable amount of GA3 (21.05 and 23.16 respectively) except for *E. hormaechei* 15a1 where none was detected in both NBRIP and Aleksandrov cultures. Zinc solubilization was not detected for all strains since no halo zone was exhibited surrounding the bacterial colonies. Siderophore production was tested positive for all strains when intense yellowish-orange halo zone appeared on plates around bacterial colonies after the 5^th^ day of incubation. EPS production was detected positive only by *E. cloacae* 38 grown on both NBRIP and Aleksandrov agar plates with the formation of mucoid-like colonies (S2 Table).

### Biosafety characteristic of *Enterobacter* spp

Blood agar was used to identify the biosafety characteristic of the *Enterobacter* spp. Haemolytic activity was not exhibited by all strains since no detectable zone of clearings (**γ**-haemolysis) by bacterial colonies on the blood agar (S1 Fig). This could indicate that the *Enterobacter* spp. may not be pathogenic for humans, thus safe for further plant bioassay studies.

### Germination assay and colonization detection of *Enterobacter* spp

Germination bioassay was carried out to evaluate the effects of *Enterobacter* spp. on the early vegetative growth stage of okra seedling. Seed biopriming using the *Enterobacter* spp. culture significantly improved (P<0.05) okra early seedling growth as compared to the uninoculated control seeds based on the length of hypocotyl and radicle, number of lateral roots and vigor index (Fig 5). The longest hypocotyl and radicle length as well as the highest number of lateral roots were recorded by seeds bioprimed with *E. hormaechei* 15a1 (16.2 cm, 8.4 cm and 13.7 respectively). Although the germination percentage remained as 100% for all treatment, the vigor index showed significant improvement for bioprimed seeds where the highest index was recorded by seeds treated with *E. hormaechei* 15a1 (2451.7) followed by *E. hormaechei* 40a (2171.7) and *E. cloacae* 38 (1998.3).

**Fig 5.**
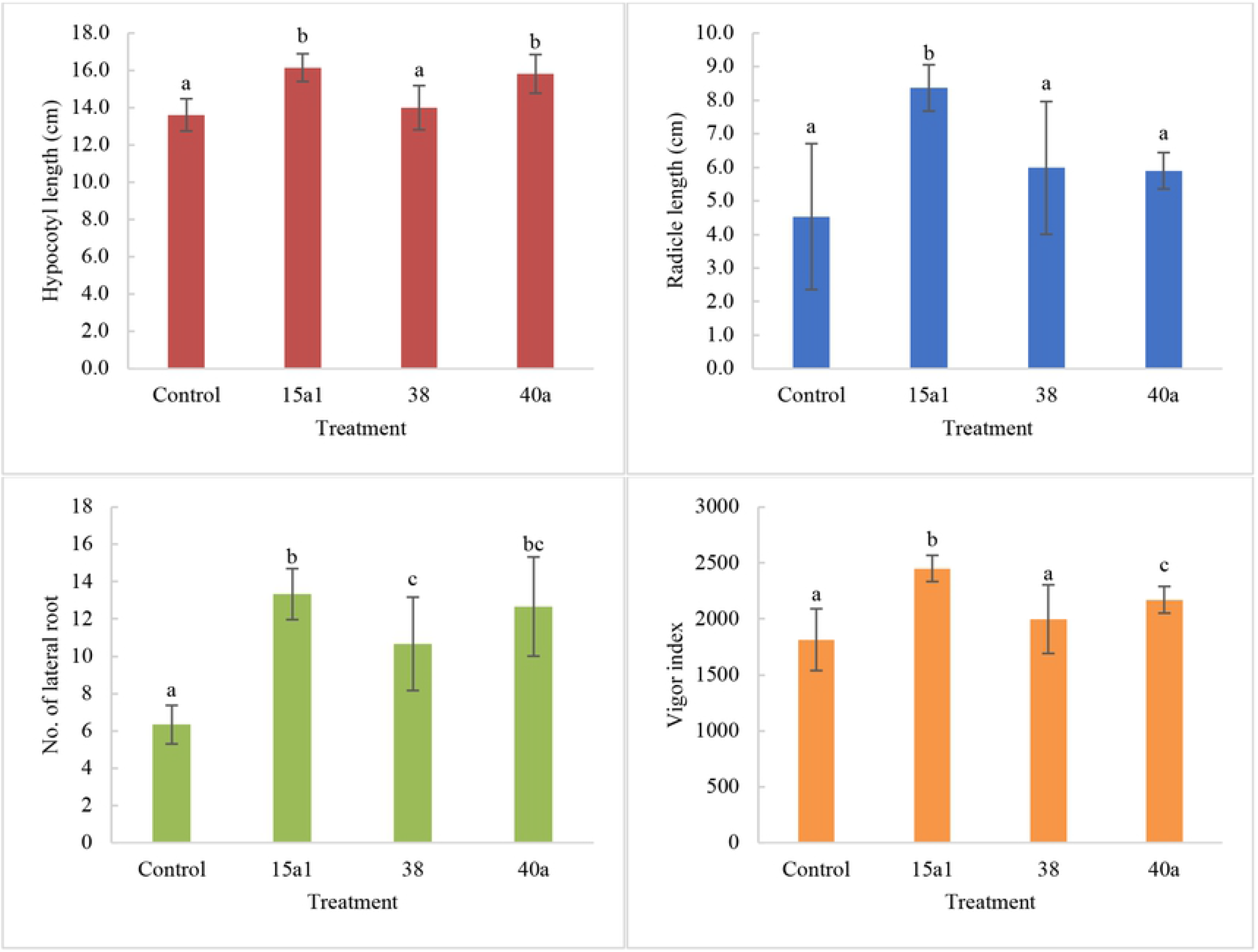
Effects of seed biopriming on germination of okra seeds in a culture tube assay under sterile tissue culture room conditions. All values are the mean value of six biological replicates alongside the standard deviation indicated by the error bar. Means followed by different letters in the same graph indicate significant differences at P<0.05.

To verify the effective colonization of *Enterobacter* spp. on okra seedling roots under sterile condition, the chemically fixated root samples were viewed using SEM. Images of root sections viewed under 3000× magnification (Fig 6) revealed that *Enterobacter* spp. successfully colonized root surface of okra seedlings where the total bacterial population for roots colonized by *E. hormaechei* 15a1, *E. cloacae* 38 and *E. hormaechei* 40a was 0.19 cells/μm^2^, 0.45 cells/μm^2^ and 0.25 cells/μm^2^ respectively.

**Fig 6.**
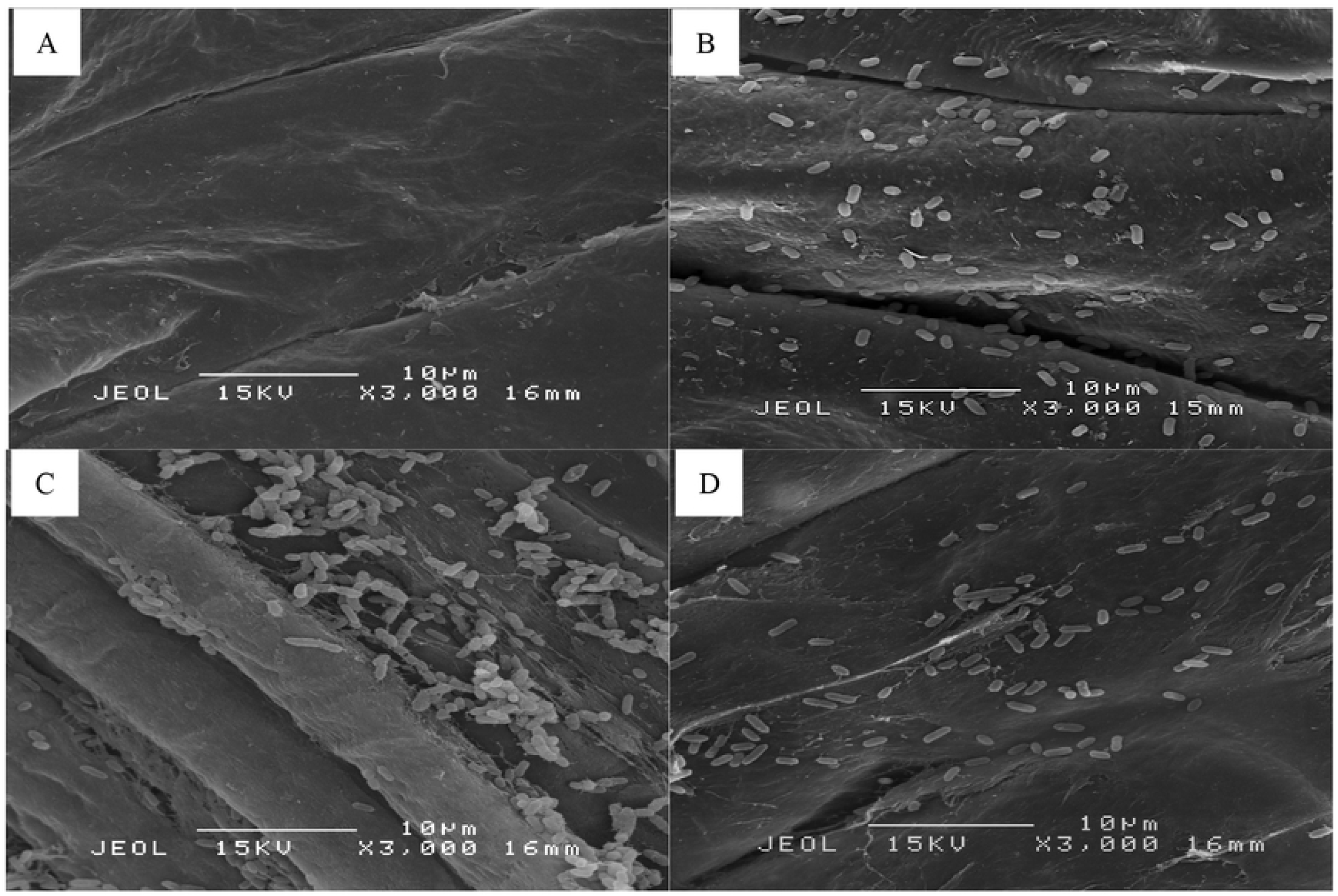
Images of root sections indicating *in situ* colonization of bacteria under controlled germination assay viewed using SEM. A: Control (uninoculated), B, C and D: Root colonization of *E. hormaechei* 15a1, *E. cloacae* 38 and *E. hormaechei* 40a respectively.

### Effects of seed biopriming with *Enterobacter* spp. on okra seedling growth

Since *Enterobacter* spp. seed biopriming gave out positive effects on okra seedling vigor index, a seedling pot experiment was carried out to evaluate the growth promoting effects under net house conditions. After 25 days of post-biopriming, seedlings were uprooted and analysed as described in the previous section. *Enterobacter* spp. seed biopriming improved early vegetative growth of okra seedlings under net house conditions. Okra seedlings treated with *Enterobacter* spp. showed a significant increase (P<0.05) in overall shoot and root length as well as plant biomass (Table 4). The shoot and root length of the bioprimed seedlings ranged from 17 to 17.4 cm and 21.4 to 24.1 cm respectively as compared to the uninoculated seedlings (14.8 and 18.8 cm respectively). Seedlings treated with *E. hormaechei* 40a was observed to show the highest fresh and dry biomass of both shoot and root plant parts.

**Table 4.** Effects of *Enterobacter* spp. on early vegetative growth of okra seedlings under net house conditions.

| Treatment | Shoot |  |  | Root |  |  |
| --- | --- | --- | --- | --- | --- | --- |
|  | Length (cm) | Fresh biomass (g) | Dry biomass (g) | Length (cm) | Fresh biomass (g) | Dry biomass (g) |
| Control (uninoculated) | 14.8±1.8a | 1.32±0.16a | 0.24±0.03a | 18.8±3.3a | 0.88±0.37a | 0.12±0.04a |
| *E. hormaechei* 15a1 | 17.4±0.6b | 1.73±0.23b | 0.28±0.04a | 22.1±2.2b | 1.76±0.41b | 0.16±0.03ab |
| *E. cloacae* 38 | 17.0±1.2b | 1.91±0.29b | 0.28±0.06a | 21.4±1.9ab | 1.94±0.26b | 0.18±0.04b |
| *E. hormaechei* 40a | 17.0±0.8b | 1.97±0.15b | 0.28±0.03a | 24.1±1.5b | 2.03±0.24b | 0.18±0.03b |

Data are mean value of six biological replicates with ± symbol denoting standard deviation. Means followed by different letters in the same column indicate significant differences at $P<0.05$ .

Leaf surface area and SPAD chlorophyll index of bioprimed seedlings were increased significantly (P<0.05) as compared to uninoculated seedlings (Table 5). Treatment using *E. hormaechei* 40a showed the highest improvement in leaf surface area (18.5 cm^2^) followed by *E. hormaechei* 15a1 (18.5 cm^2^) and *E. cloacae* 38 (15.7 cm^2^). Likewise, the SPAD chlorophyll index of the leaves of the bioprimed seedlings was increased up to 37.9. Soil viable cell of bioprimed seedlings recorded a higher CFU ranged from 6.0 × 10^7^ to 9.5 × 10^7^ CFU/g soil as compared to uninoculated seedlings (3.5 × 10^7^).

**Table 5.**
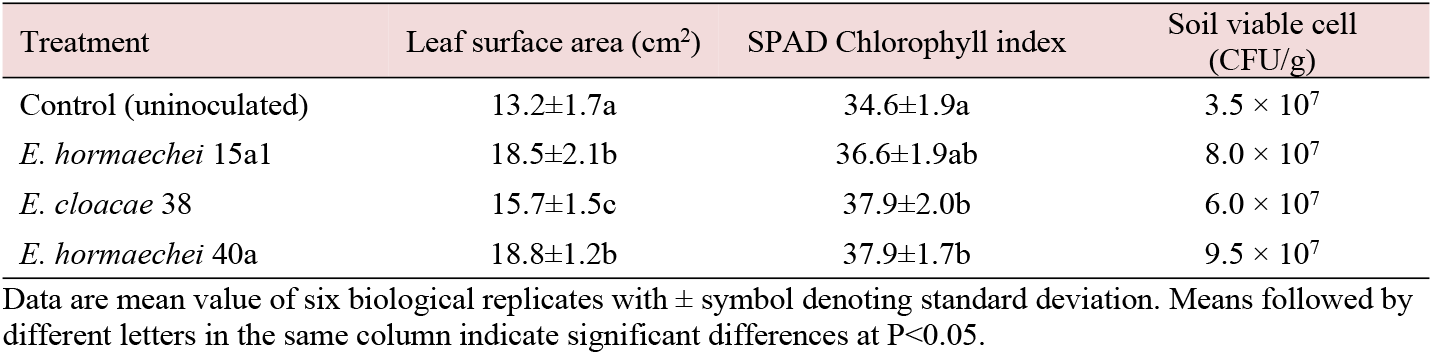
Effects of *Enterobacter* spp. on leaf surface area, SPAD chlorophyll index and soil viable cell.

| Treatment | Leaf surface area ($\text{cm}^2$) | SPAD Chlorophyll index | Soil viable cell (CFU/g) |
| --- | --- | --- | --- |
| Control (uninoculated) | 13.2±1.7a | 34.6±1.9a | $3.5 \times 10^7$ |
| *E. hormaechei* 15a1 | 18.5±2.1b | 36.6±1.9ab | $8.0 \times 10^7$ |
| *E. cloacae* 38 | 15.7±1.5c | 37.9±2.0b | $6.0 \times 10^7$ |
| *E. hormaechei* 40a | 18.8±1.2b | 37.9±1.7b | $9.5 \times 10^7$ |

Data are mean value of six biological replicates with ± symbol denoting standard deviation. Means followed by different letters in the same column indicate significant differences at $P<0.05$ .

### Effects of *Enterobacter* spp. on P and K uptake

P and K contents in plants were evaluated separately by plant sections which were leaf, stem and root along with the growing medium soil. P and K uptake of bioprimed seedlings were significantly higher (P<0.05) as compared to the uninoculated seedlings (Table 6). The range of total P content in plant sections of bioprimed seedlings was around 1.54 to 3.54 mg/g while the uninoculated seedlings were around 1.54 to 2.54 mg/g only. Treatment of *E. hormaechei* 40a consistently showed the highest total P content which included leaf (3.01 mg/g), stem (3.54 mg/g), root (1.91 mg/g) and soil (0.32 mg/g). A similar pattern was discovered in total K content of plant sections of bioprimed seedlings where the leaf (15.67 – 18.49 mg/g), stem (13.89 – 17.83 mg/g) and root (13.94 – 17.99 mg/g) were higher than uninoculated seedlings (9.79 - 12.29 mg/g). Soil available P and K were increased for bioprimed seedlings, ranging from 24.53 to 26.27 μg/g and 6.93 to 12.00 μg/g respectively over the uninoculated seedlings.

**Table 6.** Effects of *Enterobacter* spp. on P and K uptake of okra seedlings and soil available P and K under net house conditions.

| Treatment | Leaf | Stem | Root | Soil | Soil available P content (µg/g) |
| --- | --- | --- | --- | --- | --- |
|  | Total P content (mg/g) |  |  |  |  |
| Control (uninoculated) | 2.05±0.02a | 2.51±0.07a | 1.54±0.02a | 0.25±0.01a | 17.20±2.80a |
| *E. hormaechei* 15a1 | 2.90±0.05b | 3.18±0.07b | 1.60±0.02b | 0.28±0.01b | 25.47±0.92b |
| *E. cloacae* 38 | 2.63±0.09c | 2.54±0.03a | 1.54±0.03a | 0.27±0.01b | 24.53±2.05b |
| *E. hormaechei* 40a | 3.01±0.01d | 3.54±0.14c | 1.91±0.02b | 0.32±0.01c | 26.27±2.20b |
| Treatment | Total K content (mg/g) |  |  |  | Soil available K content (µg/g) |
| Control (uninoculated) | 9.79±0.08a | 11.00±0.18a | 12.29±0.08a | 0.51±0.02a | 5.20±2.23a |
| *E. hormaechei* 15a1 | 16.10±0.12b | 15.37±0.67b | 16.85±0.55b | 0.64±0.02b | 9.20±2.00b |
| *E. cloacae* 38 | 15.67±0.61b | 13.89±0.45c | 13.94±0.35c | 0.53±0.02a | 6.93±0.23a |
| *E. hormaechei* 40a | 18.49±0.49c | 17.83±0.35d | 17.99±0.14d | 0.75±0.02c | 12.00±1.83b |

Data are mean value of six biological replicates with ± symbol denoting standard deviation. Means followed by different letters in the same column indicate significant differences at P<0.05.

## Discussion

The exorbitant cost of chemical fertilizer and the limited sources of unsustainable P- and K-based mineral extraction have driven the search for alternative strategies to meet the agricultural demand for fertilizer input. The use of plant-beneficial microbes for agriculture to improve crop productivity has been taken seriously by a number of countries worldwide due to its notable efficacy. The present study was primarily designed to screen for P- and K-solubilizing bacteria and thereafter evaluated for other plant beneficial traits for the purpose of agricultural application. Besides, the root colonization ability of the strains was observed and inferred as a bacterial symbiotic mechanism to make way for available nutrients for plant nutrition. To this end, the pot experiment was applied to assess the efficiency of the strains to improve early vegetative growth of okra seedlings via seed and soil biopriming technique.

Out of 18 phenotypically different strains, 8 strains exhibited halo zone on NBRIP agar while only 3 strains showed halo zone on Aleksandrov agar. A modified NBRIP and Aleksandrov agar were incorporated in the subsequent experiment using indicator dyes to observe the pH change of agar due to P and K solubilization activity [22,23]. Since the PSI and KSI analysis of modified agar was more sensitive than the original one, therefore, the modified NBRIP and Aleksandrov agar can be regarded as more reliable indicator of P and K solubilizing activity of bacteria. Besides, the pH change can be simultaneously evaluated as an indicator of P and K solubilization capacity.

The selected strains with both P and K solubilizing ability were subjected to antagonistic interaction assay where strains were grown against each other. None of the strains demonstrated inhibitory activity, thus presumptively suitable for mixed culture experiment in quantitative estimation of P and K solubilization test. However, in the quantitative P and K solubilization assay, the mixed culture neither show synergistic effects of solubilization nor the reduction of pH of culture (P>0.05). This phenomenon could be due to the biological factors among the strains that were almost similar since they were closely related to each other (similar genus). Meanwhile in other reports, co-culture of distinctive species *Burkholderia* spp.–*Aspergillus* spp. demonstrated a synergistic effect in their phosphate solubilizing capacity [51]. Synergistic effect maybe more meaningful to be assessed when one strain with a desired specific activity is paired with another strain with distinctive beneficial traits as to complement a collective purpose [1].

The most common mechanism adopted by bacteria during P solubilization was the secretion of multiple organic acids [6]. In this study, the organic acid production of *Enterobacter* spp. was analysed using HPLC where the most notable amount of gluconic acid was detected, followed by malic, citric, succinic, acetic and GA3 acid. The intense amount of gluconic acid present in the growth media was an indicator of the active conversion from its glucose precursor via the extracellular oxidation mechanism during phosphate solubilization [52]. This could explain the acidic pH of NBRIP and Aleksandrov culture during the growth of *Enterobacter* spp. which ranged from 3.78 to 4.85. We suspected that both P and K solubilization of *Enterobacter* spp. employed the similar or partially similar pathway of remobilization since both NBRIP and Aleksandrov culture contained a spiked level of gluconic acid respectively as compared to the other organic acids produced. This, however, requires further investigation to explain the action of mechanisms involved during the P and K remobilization by the *Enterobacter* spp.

PGP trait assays revealed the ability of *Enterobacter* spp. to fix N_2_, produce IAA, GA3, siderophore and EPS. Most PGPB in the previous reports regardless of their respective genera often demonstrated more than one PGP traits [6,53,54]. N_2_-fixing ability is an important trait as potential plant bioinoculant since N_2_ is the top essential macro-nutrient required for plant growth. IAA lies in the cluster of auxins family produced by plants or certain microbes as it is important for leaf formation, embryo development and root elongation [55]. A recent report has identified an *Enterobacter* sp. that was able to secret IAA up to 5561.7 μg/mL using an optimized culture condition [56]. However, in this report, none of the strains was able to show zinc solubilization activity. Zinc solubilization is commonly exhibited by rhizobacteria including those of *Enterobacter* sp., *Pseudomonas* sp. and *Bacillus* spp. and this trait has been linked to the growth promotion of various plant varieties [6,39].

Under *in vitro* germination assay, seed biopriming with *Enterobacter* spp. improved okra early seedling growth as compared to the uninoculated control seeds based on the length of hypocotyl and radicle, number of lateral roots and vigor index. Effective root colonization of *Enterobacter* spp. was observed using SEM where a large number of bacterial cells was detected on root tips. This finding was comparable to the *in vitro* plate bioassay carried out in the previous report [6] as the seed inoculation of wheat by P solubilizing bacteria *(Enterobacter* spp. MS32) increased germination and seedling growth as well as the root volume and tips size. Similarly, seed biopriming with a series of endophytic bacteria on muskmelon seeds resulted in the promotion of of hypocotyl and radicle growth [57].

In pot experiment of okra seedling, *Enterobacter* spp. showed the potential to improve the growth of the plant as indicated by the in overall shoot and root length as well as the plant biomass. In fact, the leaf surface area and the SPAD chlorophyll index were increased significantly (P<0.05) as compared to the uninoculated seedlings. The increase of total P and K content in the bioprimed seedlings as compared to control plants could explain the reason behind the growth promotion of okra seedlings. The soluble P and K in the soil matrix were also increased providing more readily available nutrient for plant uptake. Since P and K availability is considered limiting step in plant growth and nutrition [58], this evidence suggests a significant contribution of P- and K-solubilizing *Enterobacter* spp. to promote the okra plant growth since early vegetative stage of seedling. Additionally, a recent review of biopriming techniques validated that the major microbial contributions towards enhanced plant productivity included and not limited to, N_2_ fixation; P and K remobilization; iron sequestration; microbial phytohormones, vitamins, functional enzymes, and beneficial organic acids secretion; nutrient uptake facilitation via hyphal networks; active metabolites production; seed emergence promotion; vigorous seedlings establishment [59].

## Conclusion

To the best of our knowledge, this is the first report to simultaneously focus on both P and K solubilization by *Enterobacter hormaechei* sp., highlight the plant beneficial traits and apply to pot experiment to measure the plant nutrient uptake on okra seedlings using seed biopriming technique. Specifically, increased available P and K in the soil was observed in addition to the higher P and K uptake of roots and shoots of okra seedlings. Therefore, the P and K solubilizing *Enterobacter* spp., especially *Enterobacter hormaechei* 40a might be considered as a promising candidate for P and K solubilizing biofertilizer for plant growth promotion to enhance seed germination to the subsequent early vegetative growth of seedlings and eventually the yield components. Further studies on genomic and metabolomic facets of this species might be meaningful to comprehend its broad potential in promoting crop productivity particularly the systems level response to metabolic lesions in biochemical pathways of enzymes, their physiological and ecological roles that might be appreciated in agriculture.

## Acknowledgements

This research was supported by the Putra High Impact Grant of Universiti Putra Malaysia (9659800). The authors greatly appreciate Mohamad Nor Izzalan Sohedein for the assistance provided during the plant nutrient analysis and biochemical experiments of this project.

## Supporting information

**S1 Table. Closest strains to *Enterobacter* spp. on NCBI GenBank database.**

**S1 Fig. Qualitative assay of siderophore, biosafety plates and EPS on *Enterobacter* spp.**

**S2 Fig. Organic acid profiling of NBRIP and Aleksandrov media by *Enterobacter* spp. cultures respectively using HPLC.**

